# Fitness Tradeoffs of Multidrug Efflux Pumps in *Escherichia coli* K-12 in Acid or Base, and with Aromatic Phytochemicals

**DOI:** 10.1101/2023.07.17.549369

**Authors:** Yangyang Liu, Andrew M. Van Horn, Minh T. N. Pham, Bao Ngoc N. Dinh, Rachel Chen, Slaybrina D. R. Raphael, Alejandro Paulino, Kavya Thaker, Aaryan Somadder, Chelsea C. Menke, Zachary C. Slimak, Joan L. Slonczewski

**Author notes:** Corresponding author: Joan L. Slonczewski. These two authors contributed equally (Yangyang Liu, Andrew M. Van Horn).

## Abstract

Multidrug efflux pumps are the frontline defense mechanisms of Gram-negative bacteria, yet little is known of their relative fitness tradeoffs under gut conditions such as low pH and the presence of antimicrobial food molecules. Low pH is important as it contributes to the proton-motive force (PMF) that drives most efflux pumps. We show how the PMF-dependent pumps AcrAB-TolC, MdtEF-TolC, and EmrAB-TolC undergo selection at low pH and in the presence of membrane-permeant phytochemicals. Competition assays were performed by flow cytometry of co-cultured *Escherichia coli* K-12 strains possessing or lacking a given pump complex. All three pumps showed negative selection under conditions that deplete PMF (pH 5.5 with CCCP, or at pH 8.0). At pH 5.5, selection against AcrAB-TolC was increased by aromatic acids, alcohols, and related phytochemicals such as methyl salicylate. The degree of fitness cost for AcrA was correlated with the phytochemical’s lipophilicity (logP). MdtEF-TolC and EmrAB-TolC each conferred a fitness cost at pH 5.5, but salicylate and benzoate conferred a net positive fitness contribution for the pump. Expression of pump genes was measured by digital PCR. Between pH 5.5 – 8.0, *acrA* and *emrA* were upregulated in log phase, whereas *mdtE* expression was upregulated in transition-to-stationary phase and at pH 5.5 in log phase. Methyl salicylate did not affect pump gene expression, despite selecting against AcrAB-TolC. Our results suggest that lipophilic non-acidic molecules select against a major efflux pump without positive section for others.

**IMPORTANCE:** For drugs that are administered orally, we need to understand how ingested phytochemicals modulate intrinsic drug resistance in our gut microbiome. Intrinsic drug resistance of bacteria is mediated by PMF-driven pumps that efflux many different antibiotics and cell waste products. These pumps play a key role in bacterial defense by conferring low-level resistance to antimicrobial agents at first exposure, while providing time for a pathogen to evolve resistance to higher levels of the antibiotic exposed. Nevertheless, efflux pumps confer energetic costs due to gene expression and pump energy expense. The bacterial PMF includes the transmembrane pH difference (ΔpH) which may be depleted by permeant acids and membrane disruptors. Understanding the fitness costs of efflux pumps may enable us to develop resistance breakers, that is, molecules that work together with antibiotics to potentiate their effect. We show that different pumps have distinct selection criteria, and we identified non-acidic aromatic molecules as promising candidates for drug resistance breakers.

## INTRODUCTION

The global public health threat of multidrug resistance (resistance of one strain to three or more antibiotics) (1, 2) requires us to discover new antibiotics, and to find ways of reversing or breaking resistance (3, 4). Among the various mechanisms of drug resistance, bacteria encode efflux pumps that act as a frontline defense by pumping out low levels of antibiotic (5). Efflux pumps evolved as general-purpose waste transporters that export a variety of substrates, including biosynthesis intermediates (6). These pumps support the cell’s fundamental metabolism and growth (7). However, these general transporters can also be overexpressed to enable low-level resistance that provides enough time for bacteria to evolve higher levels of resistance (8).

To study efflux pumps and identify resistance (3, 9) our lab investigates evolutionary tradeoffs between strains that possess the pump and those that do not. We are especially interested in tradeoffs associated with aromatic phytochemicals and related molecules (10–12). The fitness costs of efflux pumps might arise by various means, such as the expense of their gene expression; the cytoplasmic effects of a phytochemical; and the depletion of energy spent to export substrates. Five of the six major families of efflux pumps are powered by proton motive force (PMF) whereas one family of pumps, the ABC transporters, are powered by ATP (6).

For our investigation of fitness tradeoffs, we chose three *E. coli* efflux pumps for which knockout mutations arise during long-term evolution experiments with organic acids that incur energy stress (10, 13, 14). The most studied efflux pump in Gram-negative bacteria is AcrAB-TolC, a member of the resistance-nodulation division (RND) superfamily (6, 15). AcrAB-TolC exports diverse lipophilic substrates such as β-lactam, quinolone, and macrolide antibiotics, as wells as bile acids, polyphenols, and organic solvents (7, 16–19). Another PMF-dependent RND pump is MdtEF-TolC (20, 21). MdtEF-TolC exports many of the same substrates as AcrAB-TolC (22) but it is preferentially expressed under stationary phase and at low pH and it is essential for extreme-acid survival (19, 22, 23). The EmrAB-TolC MDR pump belongs to the major facilitator superfamily (MFS) though its tripartite structure resembles that of RND pumps (24). EmrAB-TolC exports PMF uncouplers such as dinitrophenol, CCCP, and FCCP (25) as well as colistin, quinolone, and bile acids (19, 26, 27).

We investigate the basis of energy-stress tradeoffs related to pH and PMF. *E. coli* adapts to growth at a range of external pH 5.0-9.0 (28, 29) which is typically found in the human intestinal tract (30, 31). Low external pH increases the transmembrane pH difference (ΔpH) component of the PMF available to run efflux pumps (32). The increased ΔpH (and therefore PMF) can increase the rate of drug efflux (33). At the same time, efflux pumps may compete with other PMF-intensive cell functions. For example, deletion of AcrAB-TolC increases flagellar motility which varies with proton motive force (34, 35). Low pH also amplifies uptake of membrane-permeant weak acids that can impair cell growth in various ways and can reverse antibiotic resistance (10).

Antibiotic resistance can be reversed by various phytochemicals (36) and microbial natural products, particularly aromatic molecules (36, 37). Aromatic carboxylates decrease the pump fitness contribution (11). Alternatively, cell function may be impaired by proton leak, as studied in mitochondria (38–40). Protons leak through ATP synthase and other membrane proteins (41).

Certain lipophilic molecules induce membrane leaks that deplete PMF (42). A measure of lipophilicity is logP, the octanol-water partition coefficient, which is commonly considered in assessment of pharmaceutical candidates (43). More lipophilic molecules with higher logP will tend to associate with the inner membrane. Lipophilic molecules might also affect efflux pumps by destabilizing the membrane-embedded substrate-binding pockets of pump substrates such as AcrB (44, 45).

Finally, if an upregulated drug pump is not important for survival under a given condition, such as low pH or the presence of an inhibitory phytochemical, the corresponding drug pump genes would confer a negative fitness cost to the cell and be selected against. Thus, the selective effects that phytochemicals and aromatic molecules have on drug pumps may also result from their effects on the expression of drug pump genes.

Here, we used relative fitness assays by flow cytometry co-culture to explore the fitness tradeoffs of AcrAB-TolC, MdtEF-TolC, and EmrAB-TolC in the presence of a variety of naturally occurring organic molecules with diverse chemical properties (46, 47). We aimed to distinguish the basis of their effects with respect to pH stress, PMF stress, lipophilicity (membrane solubility), and gene expression. Our findings suggest promising ways to develop resistance breakers that potentiate antibiotics by decreasing intrinsic drug resistance of pathogens.

## RESULTS

### Competition assays by co-culture with flow cytometry

The relative fitness contributions of MDR efflux pumps were measured by flow cytometry in short-term weekly competition experiments (11, 12), as described under Methods. In these experiments, a strain that carries a pump gene defect, such as Δ*acrA*::*kanR*, is co-cultured with the parent strain W3110 *ΔyhdN*::*kanR* (*acrA*^+^) for approximately 30 generations over 3 days (**Fig. 1)**. Each strain expresses a different fluorophore, YFP or CFP, which identifies cells by flow cytometry (see **Table 1** for strains). The slope of YFP/CFP log_2_ ratios provides a measure of relative fitness for the pump deletion strain. For each experimental condition, an equal number of trials use YFP label for the deletion strain and for the parent strain, respectively.

**Figure 1.**
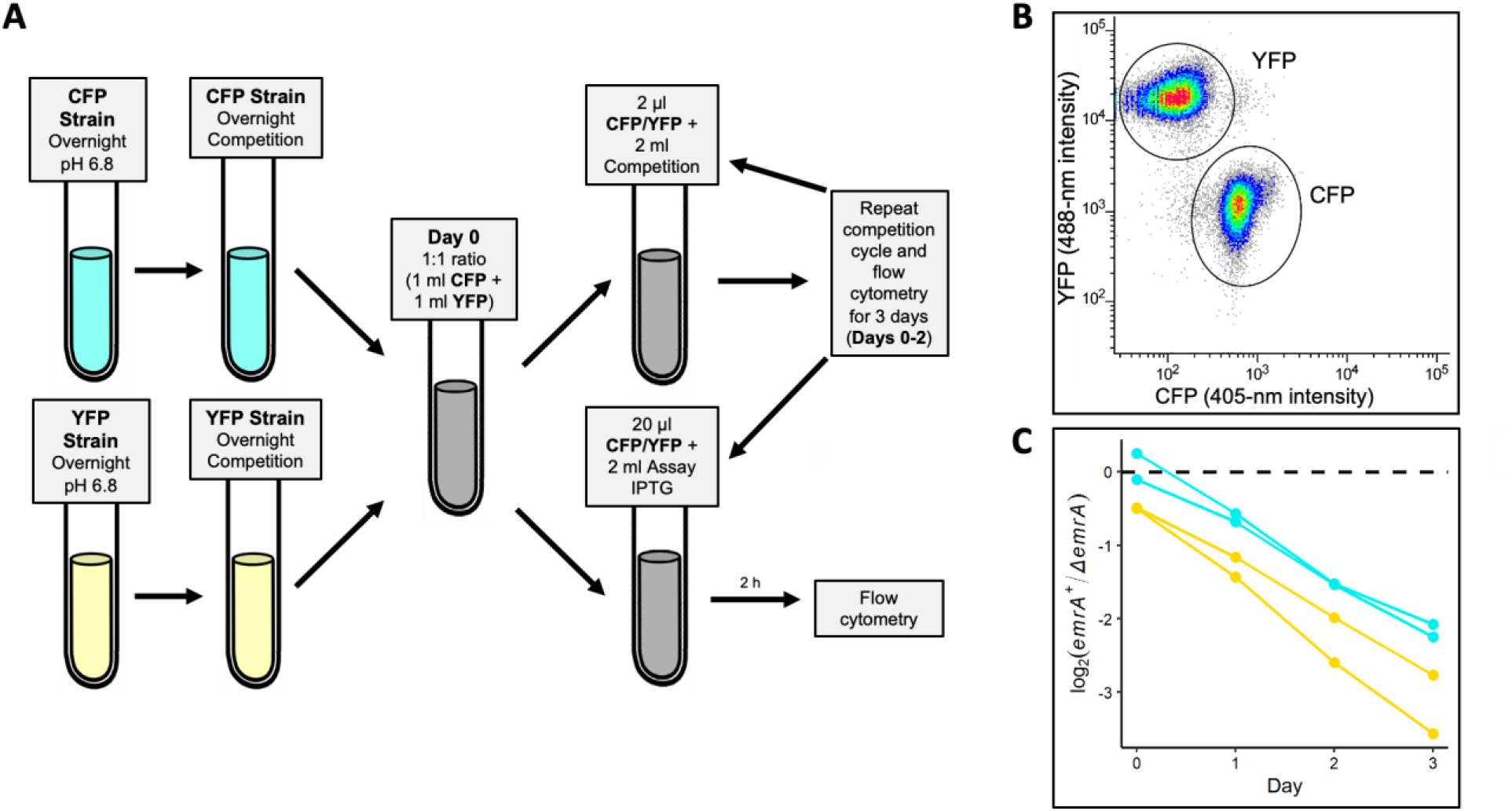
Relative fitness competition assay using flow cytometry (11). A. Daily cell co-culture was diluted 100-fold and incubated with IPTG for 2 h to induce fluorescent protein expression. B. Fluorescence intensity distribution of co-cultured YFP and CFP populations is identified by flow cytometry. The YFP-expressing population shows high-intensity emission from the 488-nm laser (Y axis) and low-intensity emission from the 405-nm laser (X axis) whereas the CFP-expression population shows the reverse pattern of emission. C. Example of a line plot displays log_2_ ratios of the pump-expressing population divided by the pump-deleted strain (*emrA^+^*/Δ*emrA*). The log_2_ population ratios are compiled for Day 0 through Day 3 for a total of ∼30 generations. The slope of the log_2_ population ratios over time (doublings per day gives the selection ratio (a measure of relative fitness) for the pump, in this case EmrAB-TolC. For each experimental condition, an equal number of trials were performed with the pump deletion linked to YFP or to CFP. For all figures, statistical tests are presented in **Table S1**.

**Table 1.** *E. coli* strains used in this study.

| Strain | Genotype | Source |
| --- | --- | --- |
| W3110 | *E. coli* K-12 F<sup>-</sup> λ<sup>-</sup> | Lab Stock (61) |
| JLS1105 | W3110 *pGFPR01*(*P<sub>BAD</sub>pHluorin bla*) | Lab Stock (28) |
| JLS1910 | W3110 Δ*galK*::*P<sub>LlacO-1</sub> cfp-bla*, Δ*yhdN*::*kanR* | Lab Stocks (11) |
| JLS1911 | W3110 Δ*galK*::*P<sub>LlacO-1</sub> yfp-bla*, Δ*yhdN*::*kanR* |  |
| JLS1826 | W3110 Δ*galK*::*P<sub>LlacO-1</sub> cfp-bla*, Δ*acrA*::*kanR* |  |
| JLS1832 | W3110 Δ*galK*::*P<sub>LlacO-1</sub> yfp-bla*, Δ*acrA*::*kanR* |  |
| JLS1834 | W3110 Δ*galK*::*P<sub>LlacO-1</sub> cfp-bla*, Δ*mdtE*::*kanR* |  |
| JLS1835 | W3110 Δ*galK*::*P<sub>LlacO-1</sub> yfp-bla*, Δ*mdtE*::*kanR* |  |
| JLS1912 | W3110 Δ*galK*::*P<sub>LlacO-1</sub> cfp-bla*, Δ*emrA*::*kanR* |  |

During passage through the digestive tract, enteric bacteria experience a wide range of pH conditions, from the stomach (pH 1.0-4.0), to the small intestine (pH 5.0-6.5) and the colon (pH 5.0-8.0) (30, 48). We compared the relative fitness contributions of efflux pumps during culture at pH values representative of the intestinal range (pH 5.5 and pH 8.0). We then tested the effects of various phytochemicals and related small molecules added to culture media at pH 5.5. For all trials, statistical tests are presented in Table S1.

### Partial PMF depletion by low-concentration CCCP or at high external pH decreases relative fitness of efflux pumps

We tested the PMF dependence of efflux-pump fitness contributions, using two different conditions that decrease PMF: growth with 10 µM CCCP at external pH 5.5, where the PMF is drained by the uncoupler; and growth at external pH 8.0, where the ΔpH is inverted and thus subtracts PMF from the electrical potential (Δψ). We tested the relative fitness for strains containing three different pump gene defects (*ΔacrA*::*kanR*, *ΔmdtE*::*kanR*, and *ΔemrA*::*kanR*) (**Fig. 2A**).

**Figure 2.**
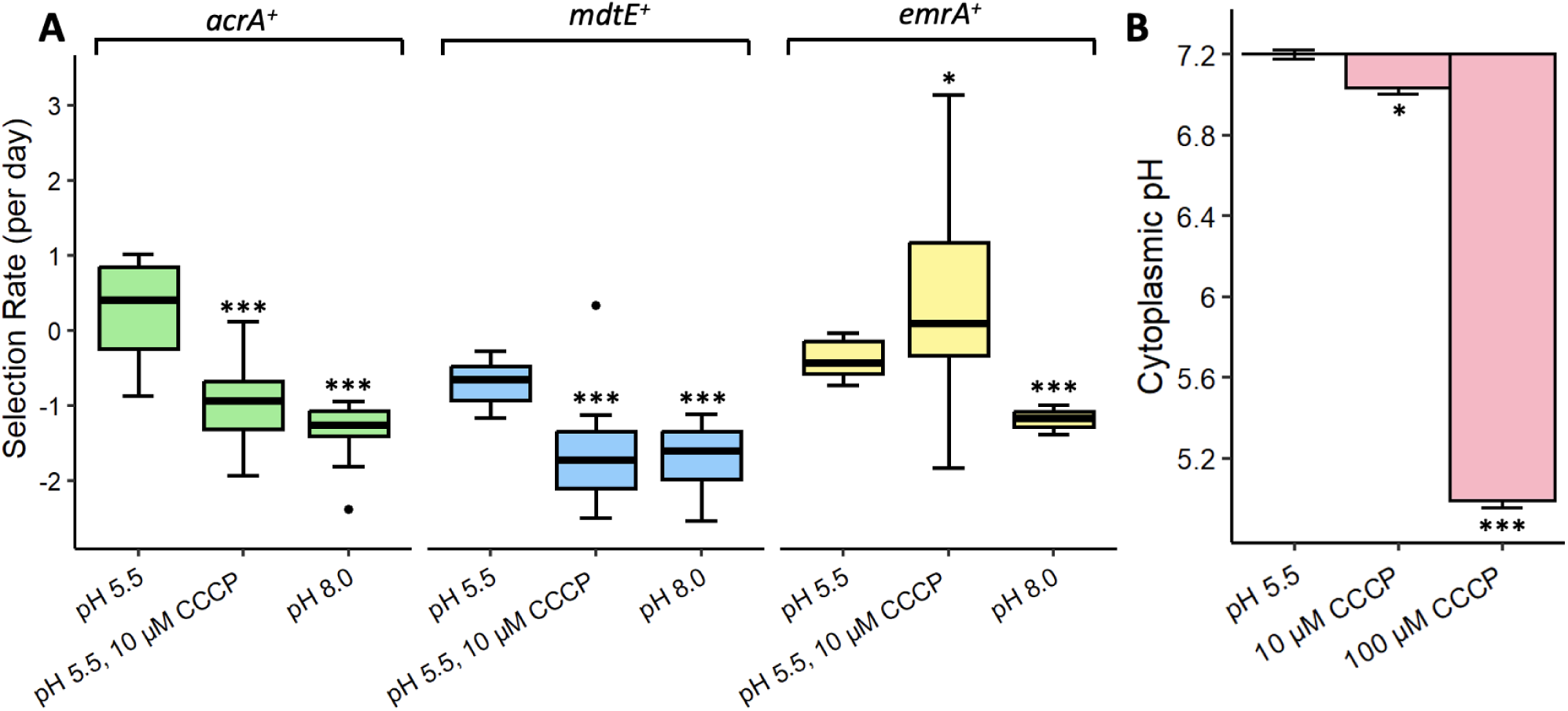
PMF depletion by CCCP or at high pH selects against *acrA^+^, mdtE^+^,* and *emrA^+^*. (A) Selection rate was calculated as shown in Figure 1. Stars indicate results of Mann-Whitney U tests for comparison of each 10 μM CCCP treatment group to the corresponding pH 5.5 control for each gene tested (*, p < 0.05; **, p < 0.01; ***, p < 0.001). (B) Change in cell pH indicates the mean difference in cytoplasmic pH of strain JLS1150 cells using pHluorin flow cytometry. Cells were cultured at external pH 5.5 with or without CCCP exposure for 5 min. Bars represent standard error. n ≥ 6 for all conditions tested. Stars indicate results of Welch’s t-tests for comparison of each CCCP treatment group to the pH 5.5 control (*, p < 0.05; **, p < 0.01; ***, p < 0.001). Statistical tests are presented in **Table S1**.

In our competition assays, a low concentration of CCCP (10 μM) at pH 5.5 selected strongly against strains expressing the pumps AcrAB-TolC or MdtEF-TolC (**Fig. 2A**). Thus, AcrAB-TolC and MdtEF-TolC each conferred a fitness cost compared to a strain deleted for a pump gene (*ΔacrA* or *ΔmdtE* respectively). The selection for EmrAB-TolC showed a wide variance, which is typical of a strong stress condition. Since CCCP is a substrate of EmrAB-TolC (25), the pump may provide a fitness advantage that partly compensates for PMF depletion.

In medium at pH 8.0, where the PMF is low, all three pumps decreased the relative fitness. The *acrA^+^*, *mdtE^+^*, and *emrA^+^* strains each produced about half as many progeny per day as their respective deletion strains (*ΔacrA*, *ΔmdtE*, and *ΔemrA*, respectively). All three pump genes showed similar fitness disparities of about one log_2_ unit between growth at pH 5.5 and pH 8.0, which were comparable to the growth disparities for *acrA^+^* and *mdtE^+^* in low-concentration CCCP at pH 5.5.

The CCCP concentration used was too small to eliminate PMF, because that condition would prevent growth. To confirm the partial depletion of PMF, we measured cytoplasmic pH in cells treated with 10 μM CCCP (**Fig. 2B**). The cytoplasmic pH was measured by use of a strain expressing the ratiometric protein fluorophore pHluorin ((28) described under Methods). A small but significant decrease in cell pH depression was observed with 10 μM CCCP (approximately - 0.2 units). A much larger pH decrease was shown in the presence of 100 μM CCCP (about −2.0 units) a concentration that prevents growth.

Overall, our results showed that partial depletion of PMF, either by growth at pH 5.5 with low concentration CCCP or by growth at pH 8.0, caused selection against the efflux pumps.

### Weak acids depress cytoplasmic pH

Membrane-permeant weak acids can depress cell pH and decrease PMF, though requiring higher concentrations than CCCP (49, 50). In addition, the growth-inhibitory anion may accumulate in the cell in proportion to the ΔpH. Both pH depression and anion accumulation may affect the fitness contribution of efflux pumps.

We compared the depression of cytoplasmic pH by various carboxylic acids and related molecules at 1 mM concentration (**Fig. 3**). The dissociation constants (pK_a_) of these molecules are in Table 2. The strongest pH depression was caused by salicylic acid (salicylate) (−0.8 pH units). At 1 mM concentration, benzoic acid, sorbic acid, and butyric acid all depressed cell pH by significant amounts. However, ferulic acid showed a slight increase in cytoplasmic pH (+0.2 units). A larger concentration of ferulic acid (10 mM) did depress cell pH (data not shown). The non-acidic molecules salicyl alcohol, vanillin, and salicylamide also caused slight increases in cytoplasmic pH, while benzyl alcohol and methyl salicylate showed no effect.

**Figure 3.**
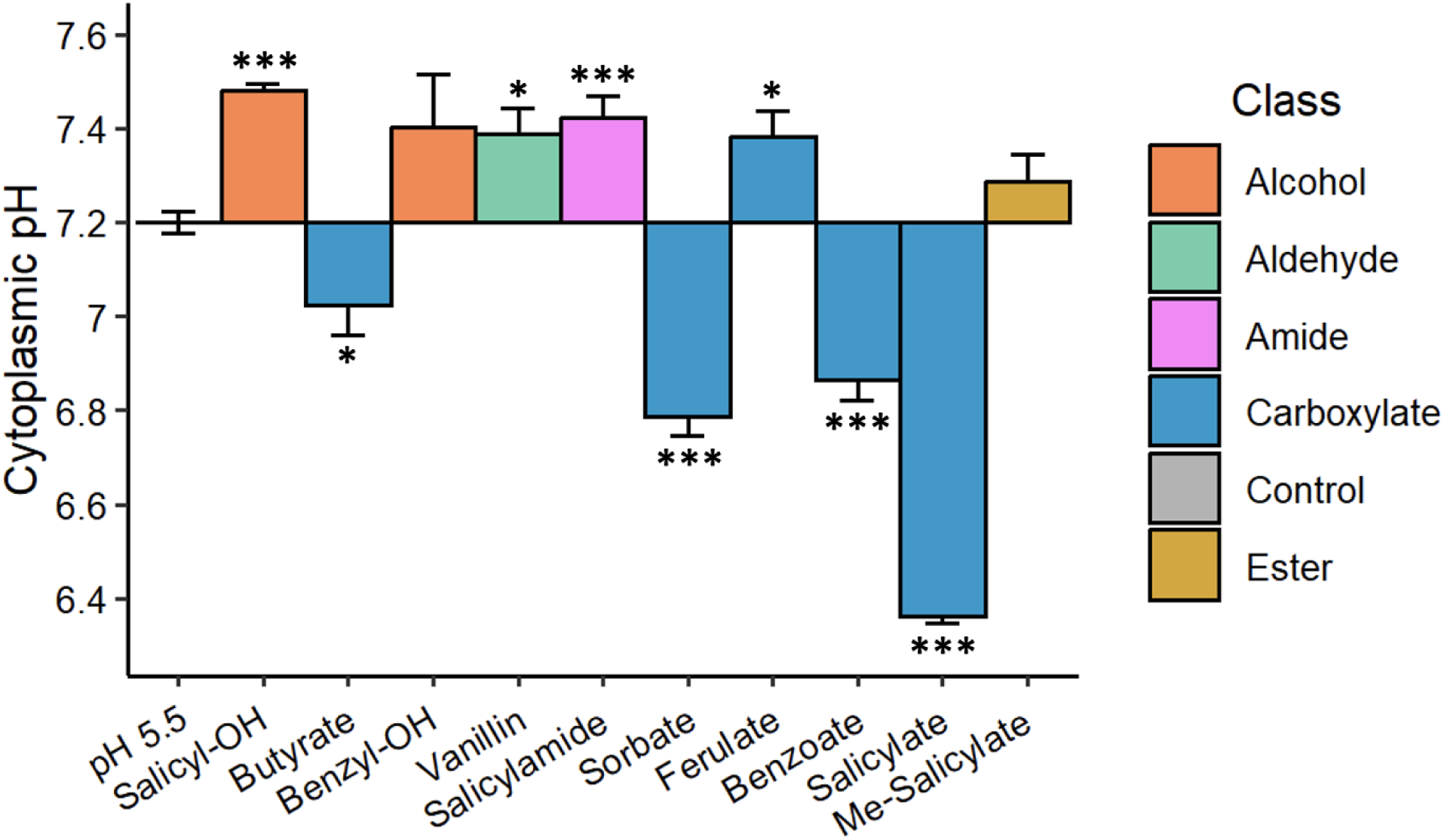
*E. coli* cell pH when exposed to various organic molecules at 1 mM. Cytoplasmic pH indicates the mean pH value after exposure to a small molecule. Values of cytoplasmic pH were determined by pHluorin fluorescence using a standard curve as described under Methods. Each culture was exposed to 1 mM of the molecule for 5 min in culture medium at pH 5.5. Colors indicate key functional groups. Bars represent standard error of the mean. Stars indicate results of Welch’s t-test for comparison of an individual molecule to the pH 5.5 control (*, p < 0.05; **, p < 0.01; ***, p < 0.001). n ≥ 6 for each of the molecules tested. Statistical tests are presented in **Table S1**.

**Table 2.** Properties for all molecules tested in FACS competition.*

| Compound | pK<sub>a</sub> | logP | MIC (mM) |
| --- | --- | --- | --- |
| Salicyl alcohol | 9.92 | 0.73 | >10 |
| Butyrate | 4.82 | 0.79 | 32 |
| Benzyl alcohol | 15.40 | 1.10 | >10 |
| Vanillin | 7.91 | 1.26 | 16 |
| Salicylamide | 8.37 | 1.28 | 16 |
| Sorbate | 4.80 | 1.33 | 4 |
| Ferulate | 4.42 | 1.51 | 16 |
| Benzoate | 4.20 | 1.87 | 4 |
| Salicylate | 3.00 | 2.24 | 2 |
| Methyl-Salicylate | 9.80 | 2.55 | >1 |
| CCCP<sup>a</sup> | 5.95 | 3.38 | 0.032 |

\*pK<sub>a</sub> and logP values sourced from open-source PubChem database. MIC values are from assays performed twice on each strain. All MIC values listed as greater than a given concentration have permitted growth at the stated concentration. CCCP = Carbonyl cyanide 3-chlorophenylhydrazone

### Lipophilic molecules select against *acrA^+^*

To investigate the mechanism of phytochemical effects on pump fitness contributions, we selected aromatic carboxylates and related small organic molecules with a range of acidity and lipophilicity (**Table 2**). For flow cytometry, strains with or without a given efflux pump were co-cultured at pH 5.5 in the presence of each selected molecule at 1 mM (**Fig. 4**). This concentration was chosen based on minimum inhibitory concentration assays showing that 1-mM concentration of each molecule allowed growth to at least OD_600_ = 0.7.

**Figure 4.**
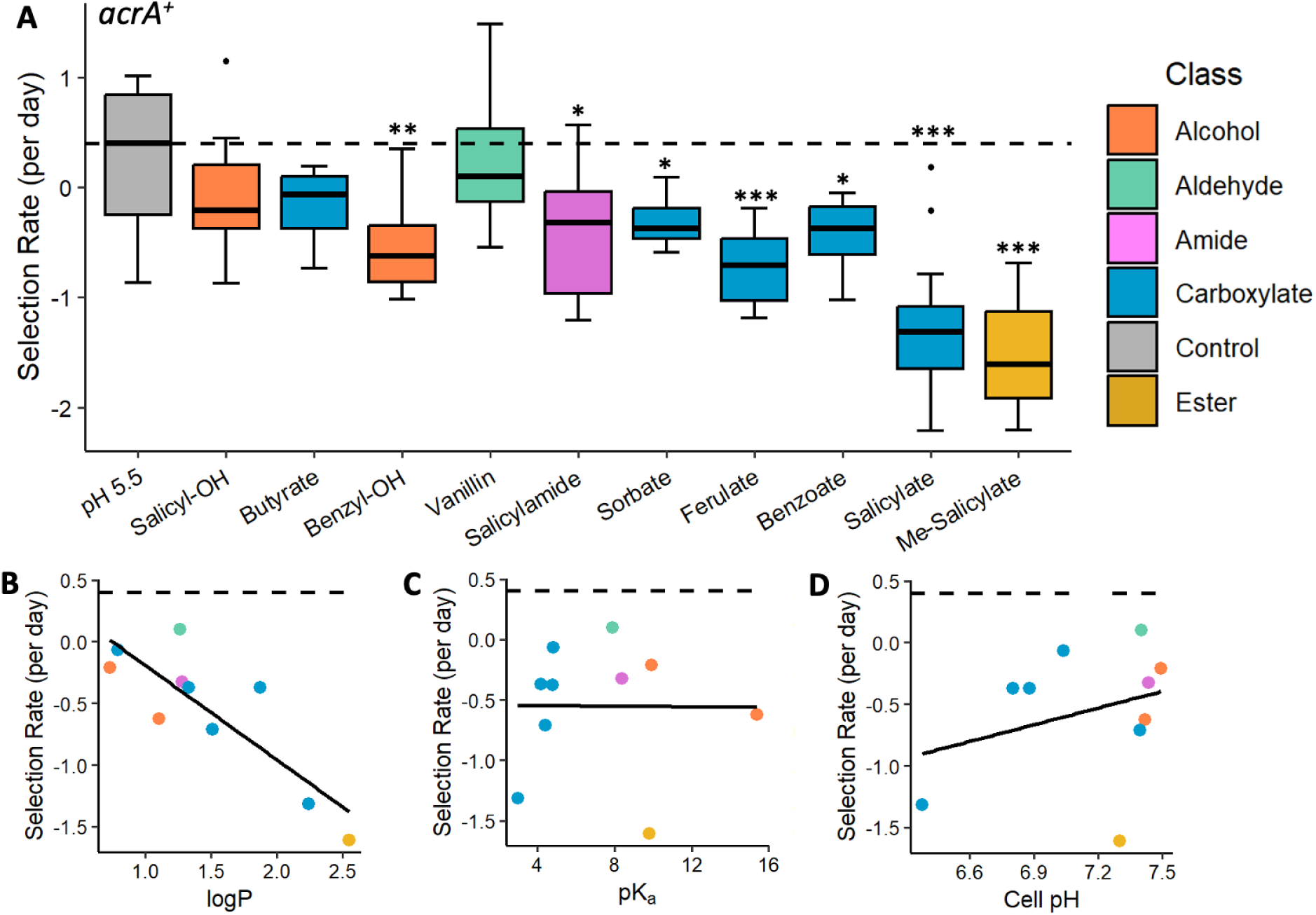
Selection for *acrA^+^* at pH 5.5. Selection rate was calculated as log_2_ *(ΔyhdN*::*kanR/ΔacrA*::*kanR*)/day. Dashed line indicates the pH 5.5 control median used as baseline for comparison. Colors indicate molecule functional groups. (**A**) Box plots of selection rates for *acrA^+^*. All molecules were tested at 1 mM. Molecules are organized from left to right in increasing order by logP. Stars indicate results of Mann-Whitney U tests for comparison of an individual molecule to the pH 5.5 control (*, p < 0.05; **, p < 0.01; ***, p < 0.001). (**B-D**) Scatter plots of median selection rates for *acrA^+^* in the presence of the molecules shown in the boxplots with Pearson least squares line. Scatter plots show relationships between selection rate for *acrA^+^*and (**B**) logP (r^2^ = 0.708), (**C**) pK_a_ (r^2^ = 7.31 x 10^-5^), and (**D**) pH change from cell pH assays after 5 minutes of 1 mM exposure to specified molecule (r^2^ = 0.095). (**A-D**) n ≥ 12 for all molecules tested. Statistical tests are presented in **Table S1**.

In the pH 5.5 control, the *acrA*^+^ strain (*ΔyhdN*::*kanR*) had a small positive selection rate (**Fig. 4A**). For further experiments, the relative fitness values for all molecules tested were nornalized to the median selection rate for the pH 5.5 control. All the tested molecules except for butyrate and salicyl alcohol selected against *acrA*^+^. Salicylate showed especially strong selection against *acrA*^+^ at a magnitude comparable to that found with CCCP or at pH 8.0 (**Fig. 2A**). But the salicylate ester, methyl salicylate, showed an equivalent size of negative selection. Thus, cell acidification was not essential for selection against *acrA^+^*.

An important factor in molecular interactions with the bacterial envelope is lipophilicity, especially for AcrAB-TolC in which the substrate binding pocket is immersed in the cell membrane (44, 45). A measure of lipophilicity commonly used to assess pharmaceutical agents is logP, representing the log_10_ of the octanol-water partition coefficient (the ratio between concentrations of the molecule in octanol versus water) (43). A strong negative correlation was observed between the *acrA^+^* selection rate and the molecule’s lipophilicity indicated by logP (r^2^ = 0.708) (**Fig. 4B**). However, no correlation was observed between *acrA*^+^ selection and pK_a_ (r^2^ = 7.31 x 10^-5^) (**Fig. 4C**) nor with cytoplasmic pH depression (r^2^ = 0.095) (**Fig. 4D**). Thus, for AcrAB-TolC the fitness tradeoff incurred by aromatic carboxylates and related molecules appeared to derive from membrane solubility rather than acidity.

### Salicylate and benzoate increase the fitness contributions of *mdtE*^+^ and *emrA^+^*

We examined the same set of structures (**Table 2**) for relative fitness assays of MdtEF-TolC. In the pH 5.5 control, *mdtE*^+^ had a negative selection rate (**Fig. 5A**). This finding suggests that MdtEF-TolC decreases bacterial fitness at pH 5.5, under our culture conditions. Compared to *acrA^+^* (**Fig. 2A**), *mdtE^+^* incurs about one log_2_ unit loss of relative fitness.

**Figure 5.**
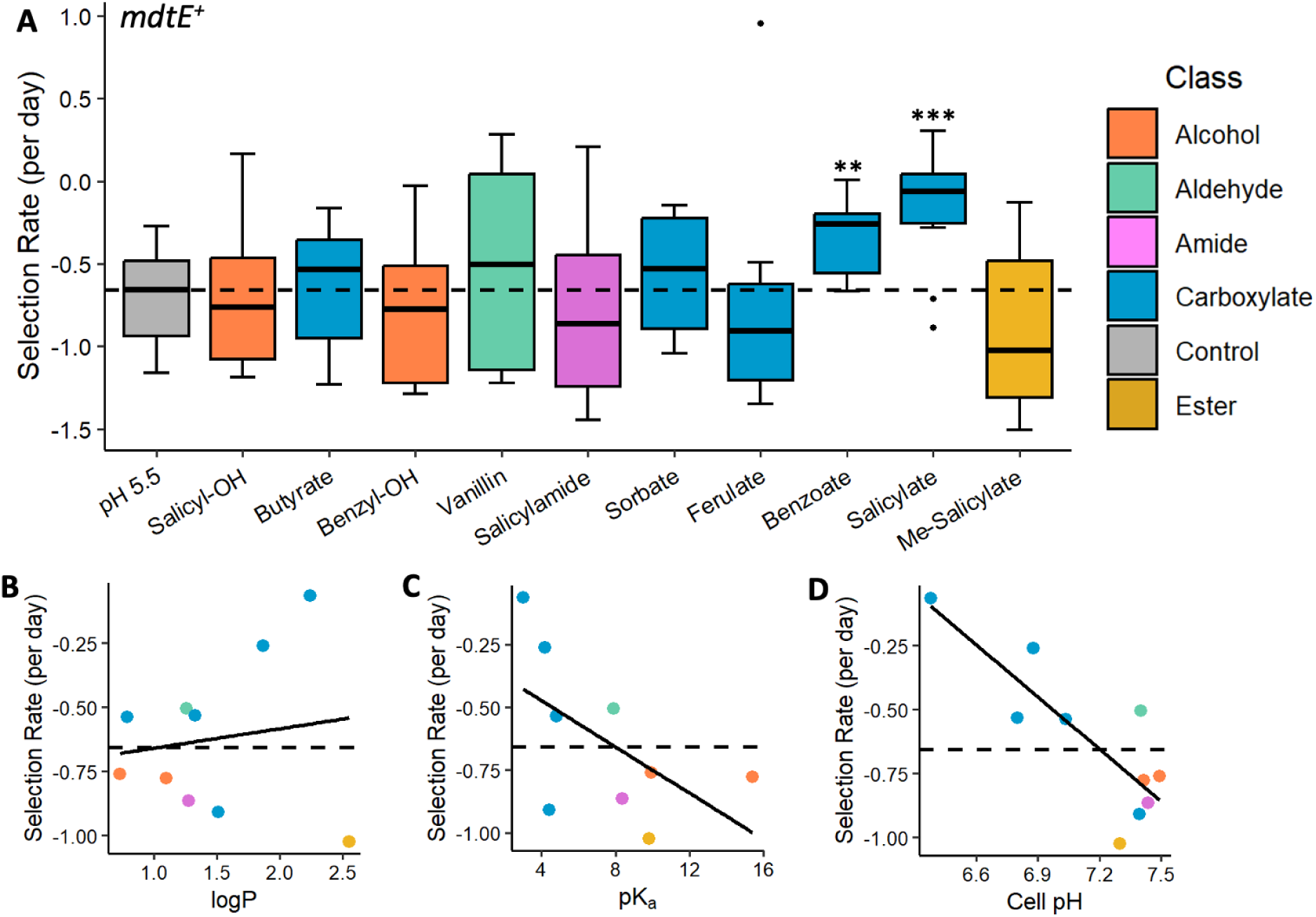
Selection for *mdtE^+^* at pH 5.5. Selection rate was calculated as log_2_ *(ΔyhdN*::*kanR/ΔmdtE*::*kanR*)/day. Dashed line indicates the pH 5.5 control median used as baseline for comparison. Colors indicate molecule functional groups. (**A**) Box plots of selection rates for *mdtE^+^*. All molecules were tested at 1 mM. Molecules are organized from left to right in increasing order by logP. Stars indicate results of Mann-Whitney U tests for comparison of an individual molecule to the pH 5.5 control (*, p < 0.05; **, p < 0.01; ***, p < 0.001). (**B-D**) Scatter plots of median selection rates for *mdtE^+^* in the presence of the molecules shown in the boxplots with Pearson least squares line. Scatter plots show relationships between selection rate for *mdtE^+^* and (**B**) logP (r^2^ = 0.022), (**C**) pK_a_ (r^2^ = 0.335), and (**D**) pH change from cell pH assays after 5 minutes of 1 mM exposure to specified molecule (r^2^ = 0.691). (**A-D**) n ≥ 12 for all molecules tested. Statistical tests are presented in **Table S1**.

Out of all molecules tested, only benzoate and salicylate significantly affected the selection rate for *mdtE*^+^. Each of these aromatic carboxylates partly reversed the negative fitness contribution of *mdtE*^+^ at pH 5.5. Sorbate and butyrate also showed a median selection rate above that of the pH 5.5 control, although the wide variance eliminated significance.

The *mdtE* selection rate showed a negative correlation with the molecule’s pK_a_ (r^2^ = 0.335) (**Fig. 5C**). Moreover, a strong negative correlation is observed (r^2^ = 0.691) between the selection rate and the molecule’s ability to depress cytoplasmic pH (**Fig. 3**, **Fig. 5D**). These results suggest that molecules that acidify the cytoplasm select for MdtEF-TolC. Unlike *acrA^+^*, no correlation was observed for *mdtE*^+^ selection with logP (r^2^ = 0.022) (**Fig. 5B**).

Similar to *mdtE^+^*, *emrA*^+^ had a negative selection rate at pH 5.5 (**Fig. 6A**), but benzoate and salicylate increased relative fitness. Salicylamide selected against *emrA^+^*. Overall, like *mdtE^+^*, the *emrA^+^* selection rate showed moderate negative correlation between relative fitness and molecule pK_a_ (r^2^ = 0.236) (**Fig. 6C**) and a strong negative correlation with the change in cytoplasmic pH (r^2^ = 0.736) (**Fig. 6D**). Only a small correlation was observed for the *emrA^+^* selection rate with logP (r^2^ = 0.127) (**Fig. 6B**). Overall, both MdtEF-TolC and EmrAB-TolC showed a positive fitness contribution in the presence of the membrane-permeant aromatic acids salicylate and benzoate.

**Figure 6.**
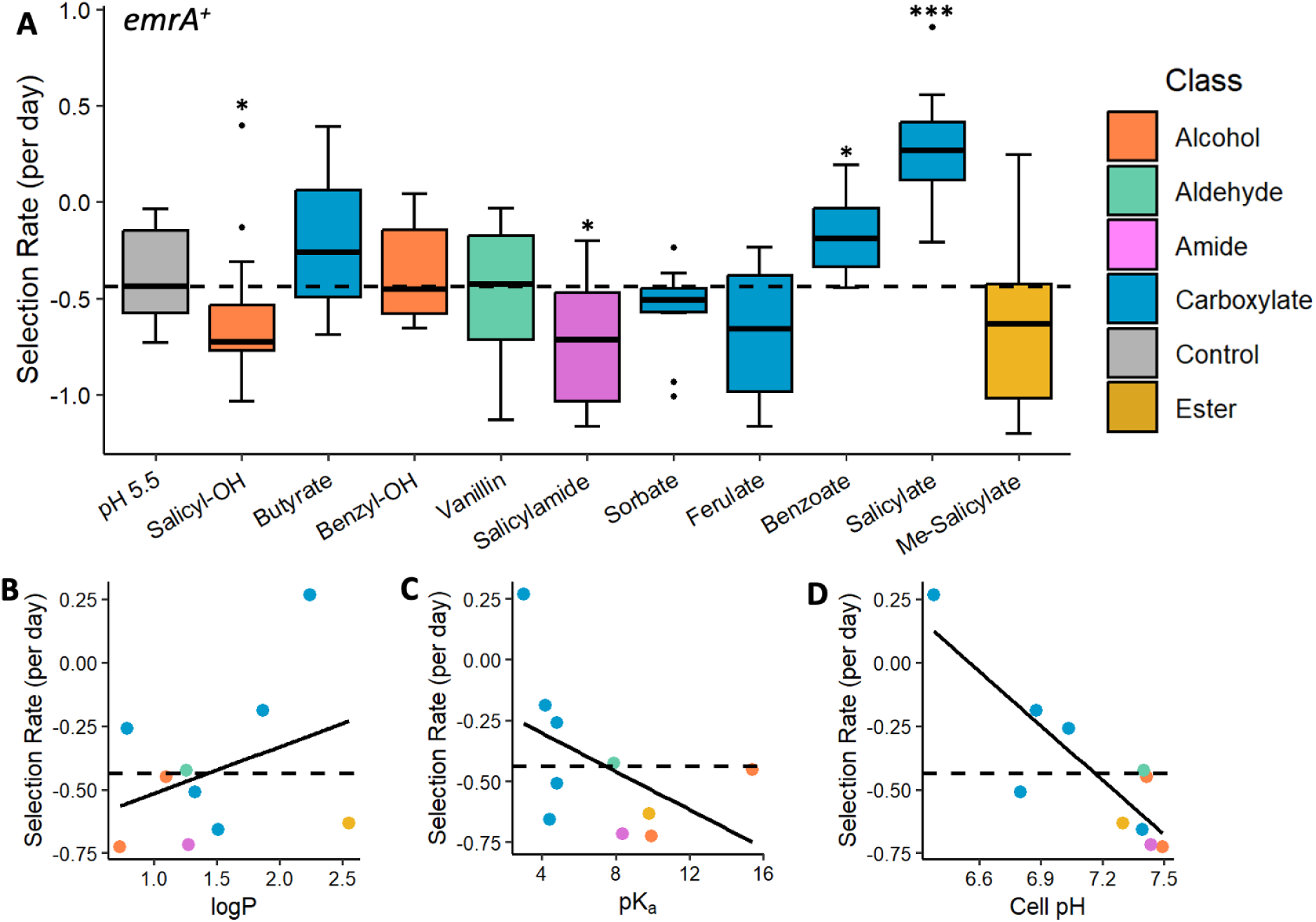
Selection for *emrA^+^* at pH 5.5. Selection rate was calculated as log_2_ *(ΔyhdN*::*kanR/ΔemrA*::*kanR*)/day. Dashed line indicates the pH 5.5 control median used as baseline for comparison. Colors indicate functional groups. (**A**) Box plots of selection rates for *emrA^+^*. All molecules were tested at 1 mM. Molecules are organized from left to right in increasing order by logP. Stars indicate results of Mann-Whitney U tests for comparison of an individual molecule to the pH 5.5 control (*, p < 0.05; **, p < 0.01; ***, p < 0.001). (**B-D**) Scatter plots of median selection rates for *emrA^+^* in the presence of the molecules shown in the boxplots with Pearson least squares line. Scatter plots show relationships between selection rate for *emrA^+^*and (**B**) logP (r^2^ = 0.127), (**C**) pK_a_ (r^2^ = 0.236), and (**D**) pH change from cell pH assays after 5 minutes of 1 mM exposure to specified molecule (r^2^ = 0.736). (**A-D**) n ≥ 12 for all molecules tested. Statistical tests are presented in **Table S1**.

### Expression profiles of pump genes vary with growth phase and pH

The fitness tradeoffs for efflux pumps may depend on the energy costs of gene expression, which varies across the phases of growth (51). Growth phase is relevant because in our co-culture experiments, the daily cycle of serial dilutions runs through lag phase, log phase, and stationary phase.

We quantified the expression of genes representing each of the three efflux pumps (*acrA*, *mdtE*, and *emrA*) by isolating RNA transcripts from cultures at three pH levels (**Fig. 7**). The cultures were each harvested during log phase (OD_600_ = 0.1-0.2) and during the transition to stationary phase (OD_600_ = 0.4 – 0.7). The OD_600_ values and time of harvest of individual replicates are presented in Table S2. Transcript levels were measured using dPCR multiplex assays (see Methods). The Taqman probe sets and fluors are presented in Table S3. Statistical tests are presented in Table S1.

**Figure 7.**
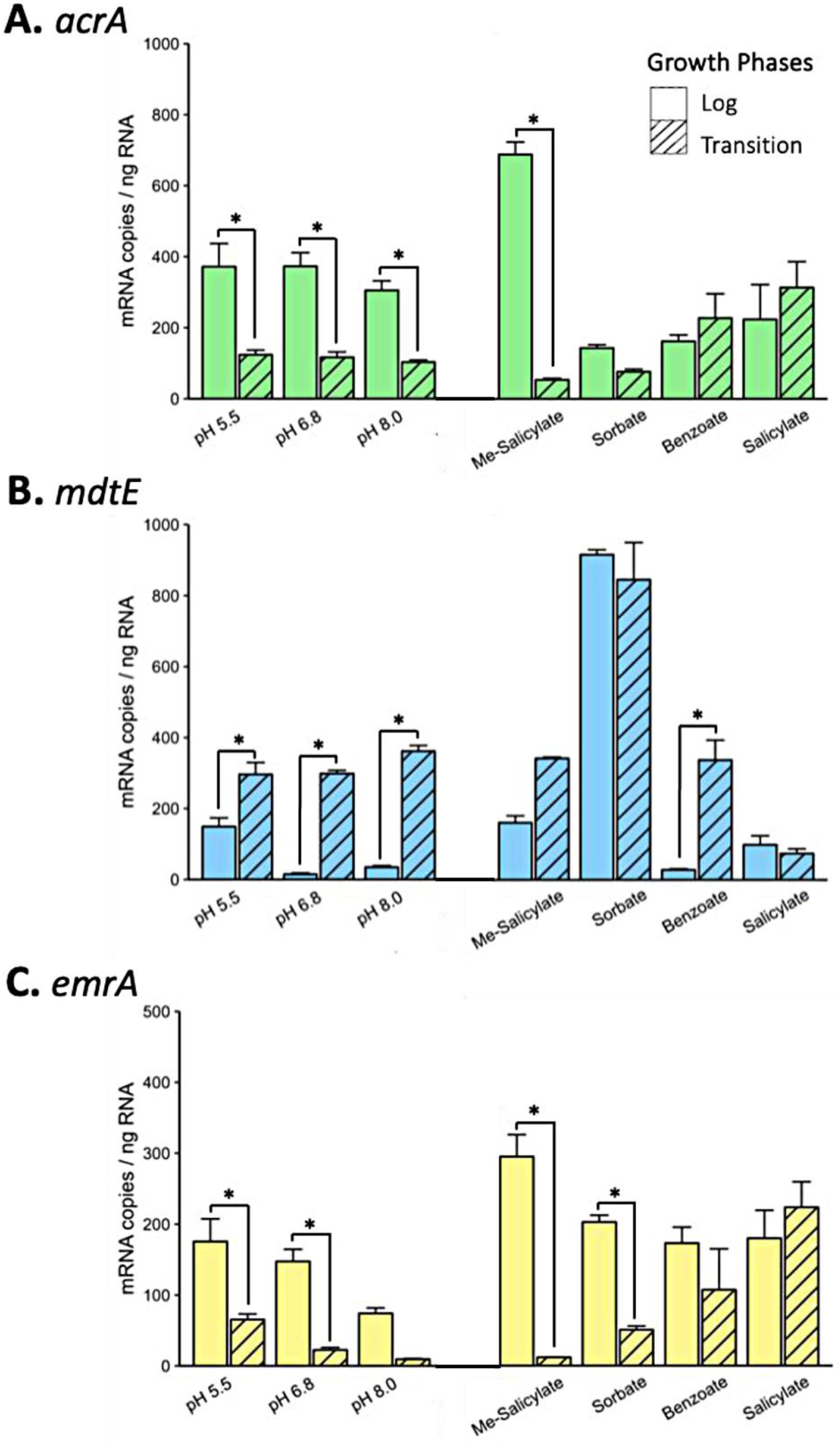
Gene expression for *acrA^+^*, *mdtE^+^*, and *emrA^+^* under various molecule conditions at pH 5.5 determined by Multiplex dPCR assay. *E. coli* K-12 W3110 was cultured to log phase and transition-to-stationary phase in LBK buffered at pH 5.5, at pH 6.8, and at pH 8.0 (Table S2). The dPCR probe sets used were for (A) *acrA*, (B) *mdtE*, (C) *emrA* (Table S3). For each probe set, the mRNA expression levels were normalized by dividing the transcript concentrations measured in dPCR (copies/μl) by the total RNA concentrations (ng/μl) of the samples used. Colors represent MDR pump genes and bar patterns represent growth phases. Error bars represent 1 SEM. Numbers along the x-axis correspond to media pH. All molecules were tested at 1 mM concentration. n = 3 for each combination of conditions (for example, *acrA*, pH 5.5, log phase). Two-way ANOVA and post-hoc least squares difference with the Benjamini-Hochberg correction were used to compare expression levels of individual efflux pump genes across all tested conditions. Statistical tests are presented in **Table S1**.

The *acrA* and *emrA* genes showed higher expression in log phase than transition phase. Across moderate pH levels (5.5 – 8.0), *emrA* expression levels were approximately half those of *acrA* expression. The *acrA* expression did not show pH dependence, but in log phase *emrA* showed higher expression at pH 5.5 than at pH 8.0. By contrast, *mdtE* showed higher expression in transition phase than in log phase, consistent with the literature (51). In log phase, *mdtE* was upregulated strongly at pH 5.5, consistent with its inclusion in the Gad acid fitness island (52) and its evidence for acid-adapted substrate binding (22). Thus, the two pumps that showed a positive fitness contribution in the presence of salicylate or benzoate (MdtEF-TolC and EmrAB-TolC) also showed upregulation in log phase at low external pH.

We also tested the expression effects of selected carboxylates and methyl salicylate, during growth at pH 5.5 (**Fig. 7**). Methyl salicylate increased the log-phase expression of *acrA*, but had marginal effets on *mdtE* and *emrA*. The inclusion of sorbate, benzoate or salicylate slowed the growth rates (Table S2) and in some cases narrowed the difference in expression between log and transition phases. In general, no pattern of gene expression appears to explain the selection effects reported in **Figures 4-6**, especially the large negative fitness contributions of salicylate and methyl salicylate for *acrA^+^* (**Fig. 4**).

## DISCUSSION

Efflux pumps serve bacteria with multiple functions for waste excretion and defense. In natural microbiomes, including those of the digestive tract, the drug resistance mediated by efflux pumps might be decreased by the presence of naturally occurring phytochemicals that act as resistance breakers (3, 4, 9). However, the basis of phytochemical resistance breakers is poorly understood.

Our results point to some common physical-chemical principles associated with molecules that select against *E. coli* strains with multidrug efflux pumps. We showed that partial depletion of PMF, by growth with CCCP or at high pH, incurs strong selection against all three efflux pumps tested (**Fig. 2**). This negative selection makes sense given the load on PMF incurred by these pumps.

Membrane-permeant acids with partial effects on PMF also selected against the AcrAB-TolC efflux pump (**Fig. 4**). But overall, neither PMF nor cytoplasmic pH depression explain the *acrA^+^* selection rates observed in the presence of various phytochemicals including non-acidic molecules such as methyl salicylate and salicylamide. However, the *acrA^+^*selection rates in the presence of the molecules tested were highly correlated with the lipophilicities of these compounds (**Fig. 4B**). Lipophilicity determines a molecule’s ability to integrate into the bacterial inner membrane, where it can interact with the membrane-embedded substrate-binding domain of AcrB in the AcrAB-TolC pump (44, 45). Such a mechanism might explain the selective effects of lipophilic non-acids such as methyl salicylate (53, 54). AcrAB-TolC is understood to primarily efflux substrates from the periplasm that dissolve in the inner membrane (55, 56).

Unlike AcrA, MdtE incurred strongly negative selection at pH 5.5 in the control medium and in the presence of most tested compounds (**Fig. 5A**). In our gene expression assay, *mdtE* was upregulated in the transition to stationary phase, a result consistent with the literature (51). This stationary-phase upregulation of MdtEF-TolC is advantageous for cells under anaerobiosis or under extreme-acid exposure (11). The *mdtE* gene has a positive fitness contribution only when the growth cycle includes a period of extreme acid (pH 2.0) comparable to stomach passage. This observation is consistent with the inclusion of *mdtEF* in the Gad acid fitness island (*slp-gadX*), which is one of the most effective mechanisms of acid resistance employed by *E. coli* (57, 58). In our competition experiments, *mdtE*^+^ showed enhanced selection rate in the presence of salicylate or benzoate, permeant acids that acidify the cytoplasm. This could result from cytoplasmic pH depression (**Fig. 3**) which is part of the mechanism of acid-stress Gad induction (59).

The effects of phytochemicals on selection rates for *mdtE*^+^ and *emrA^+^*(**Figs. 5 and 6**) were generally small. The relatively low expression of EmrAB-TolC suggests that deletion of the major pump AcrAB-TolC would be required to see larger effects associated with EmrAB-TolC (51). EmrAB-TolC did increase relative fitness in the presence of CCCP, although with a wide variance. Uncouplers such as CCCP and dinitrophenol are substrates and inducers for EmrAB-TolC (27, 60).

Overall, diverse selection criteria were observed among *acrA^+^*, *mdtE*^+^, and *emrA^+^*. The effect of lipophilic non-acids on AcrAB-TolC selection appears particularly interesting. Lipophilic aromatic esters and other non-acidic molecules could offer a promising class of potential candidates for antibiotic resistance breakers. Future investigation might reveal aromatic esters and other lipophilic molecules that act as resistance breakers for other pumps besides AcrAB-TolC.

## MATERIALS AND METHODS

### Strains and Media

The experimental strains used were derived from parental strain *E. coli* K-12 W3110 (10, 61). All recombinant strains used in flow cytometry competitions contained alleles for cyan fluorescent protein (CFP) or yellow fluorescent protein (YFP) on an IPTG-inducible lactose promoter (P_L*lac*O-1_) (10, 11, 62) (Table 1). The strain used for cell pH depression assays contained a pH-dependent fluorophore (pHluorin) (63) on an arabinose-inducible promoter (P_BAD_) in plasmid pGFPR01 (28). pHluorin shows excitation peaks at 410 nm and 470 nm that depend ratiometrically on cytoplasmic pH (63). Lab stock *E. coli* K-12 W3110 (64) was used for digital polymerase chain reaction assays.

Growth media was LBK broth (10 g/l tryptone, 5 g/l yeast extract, 7.45 g/l potassium chloride) buffered at pH 5.5 with 100 mM 2-(*N*-morpholino)ethanesulfonic acid (MES; pK_a_ = 6.15), pH 6.8 with piperazine-N,N′-bis(2-ethanesulfonic acid) (PIPES; pK_a_ = 6.8), or pH 8.0 with [4-(2-hydroxyethyl)-1-piperazineethanesulfonic acid] (HEPES; pK_a_ = 7.55). The pH was adjusted with NaOH or HCl (11). All media materials and reagents were purchased from Thermofisher unless stated otherwise: CCCP (Millipore Sigma); MES and methyl-salicylate (ACROS Organics); sodium ferulate (Selleck Chemical); and salicyl alcohol (Alfa Aesar).

### Minimum Inhibitory Concentration Assays

Strains were cultured for 22-24 h in LBK pH 5.5 buffered with 0.1 M MES, with aeration, prior to minimum inhibitory concentration (MIC) assay. 96-well plates were prepared by two-fold serial dilution with molecules of interest. Medium used in plates was LBK 0.1 M MES pH 5.5. 1 μl of overnight culture was added to each well that contained 200 μl media (1:200). OD_600_ absorbances were measured via Molecular Devices Versa *max* Tunable Microplate Reader every 15 min over 22 h (96 total reads). Plates were incubated at 37°C and shaken between reads. After 22 h, wells without growth observed were scored at or above the MIC.

### Flow Cytometry Competition Assays

Flow cytometry was used to measure the relative fitness of MDR efflux pumps in short-term weekly competition experiments across ∼30 generations (11, 12) (**Fig 1**). All cultures were incubated at 37°C with rotation. For overnight cultures, two days prior to flow cytometry (Day -2), strains were cultured individually for 22-24 h in 2 mL LBK 0.1 M PIPES pH 6.8. The next day (Day -1), cultures underwent 1000-fold dilution (2 μl into 2 ml) into weekly competition growth media. Competition media was LBK buffered at either pH 5.5 or 8.0 containing a molecule of interest at 1 mM unless stated otherwise. On Day 0, 1 ml of the pump knockout strain (*ΔacrA*::*kanR*, *ΔmdtE*::*kanR*, or *ΔemrA*::*kanR*) was mixed (1:1) with 1 ml of the control *ΔyhdN*::*kanR* containing the opposite fluorophore. Co-cultures then underwent 1000-fold dilution (2 μl into 2 ml) into fresh competition growth media prior to incubation for 22-24 h. On Days 0-2, overnight cultures were serially diluted 1000-fold (2 μl into 2 ml) into fresh competition growth media prior to incubation for 22-24 h.

For daily flow cytometry, overnight coculture were diluted 100-fold (20 μl into 2 ml) into LBK pH 6.8 0.1 M PIPES containing 1 mM isopropyl β-D-1 thiogalactopyranoside (IPTG) to induce fluorophore expression (11, 12)(65, 66). After 2 h incubation, cultures were diluted 20-fold into 1X phosphate-buffered saline (PBS). The cell suspensions were sampled with a BD FACSMelody Cell Sorter with a violet laser (405 nm) with a 528/45 filter for CFP, and a blue laser (488 nm) with a 545/20 filter for YFP emission. The proportions of processed events were >95%. PBS dilution ratio was adjusted to lower observed event rate below 10,000. Two technical replicates (each 50,000 events) were recorded, and YFP/CFP ratios were averaged. This process of recording CFP and YFP percentages was repeated for Days 0-3 of flow cytometry.

For each experimental condition tested, the selection rate for the gene tested was calculated with:

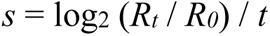

where *R_t_* / *R_0_* represents the ratio of the control strain *ΔyhdN* versus the knockout strain of interest (for example, *acrA^+^* / *ΔacrA*); and *t* represents time in days with ∼10 generation per day (65, 66). This calculation provides a biological indication of co-cultured genetic variants’ population distribution over time. For example, a selection rate of −1 indicates that there was a two-fold selection against the MDR pump-of-interest per day.

### Cytoplasmic pH Assays

Cytoplasmic pH was measured by flow cytometry of strain JLS1105 with the plasmid pGFPR01 expressing the ratiometric fluorophore pHluorin (28, 63). One day prior to flow cytometry, strain JLS1105 was cultured in 2 ml LBK-MES pH 5.5 and 100 μg/mL ampicillin at 37°C with rotation for 16-20 hours. 2 μl of overnight culture was diluted (1:1000) into 2 ml pH 5.5 0.1 M MES + 50 μg/mL Ampicillin + 0.2% L-Arabinose LBK. L-Arabinose was used to induce pHluorin expression from a pBAD promoter (28). The cultures were then incubated for 4.0-4.5 h.

After incubation, cultures were diluted ten-fold (100 μl into 1 ml) with stress growth media (pH 5.5 0.1 M MES LBK with the addition of a molecule of interest). Plain pH 5.5 0.1 M MES LBK was run at the beginning and end of all sampling sessions to ensure no significant differences overtime. 5 min after dilution, cell suspensions were sampled through BD FACSMelody. The emission intensity ratio was obtained between the 405-nm excitation (high pH) and the 488-nm excitation (low pH). 50,000 events were recorded for each trial. The processed events were always more than 95% and events rates were no more than 10,000 events/second for all samples recorded. A minimum of six biological replicates were performed for each chemical at each concentration.

Data were analyzed by converting raw fcs files to csv files using the bioconda fcsparser package. For each emission ratio, the mean, median, standard deviation, and standard error were calculated by a custom script with R and Python 3. In a given trial, if fewer than 3,000 cells were counted at 488-nm excitation, the trial was excluded from the dataset to avoid noise signal from dead or non-fluorescent cells. Mean ratios were then converted to a pH value using a standard curve for pH as a function of emission ratio. The standard curve was obtained using 40 mM sodium salicylate to equilibrate cytoplasmic and external pH over the range of pH 5.5-8.0.

### Digital Polymerase Chain Reaction (dPCR) Expression Assays

One day before harvest, *E. coli* K-12 W3110 (61) were cultured in LBK 0.1 M MES pH 5.5 containing the molecule of interest for 22-24 h at 37°C with rotation for 22 h. On the day of harvest, the final optical density of *E. coli* cultures was measured with the SpectraMax Plus384 MicroPlate reader. Cultures were diluted 1000-fold (2 μl into 2 ml) into fresh medium. The bacteria were cultured to late log phase (OD_600_ = 0.1-0.2) or transition to stationary phase (OD_600_ = 0.4-0.7).

For RNA extraction, 700 μl of each culture at log phase and 500 μl of each culture at transition-to-stationary phase (transition phase)was fixed using Bacterial RNAProtect (QIAGEN – Germantown, MD). RNA was extracted from the fixed culture pellets using *E. coli*-specific adaptations of the RNA extraction protocols in the RNAProtect Bacteria Reagent Handbook (QIAGEN). Following RNA extraction, RNA quality and quantity was measured with Nanodrop One Spectrophotometer (Thermo Fisher Scientific – Waltham, MA). Samples with no detectable chemical contaminants were selected for gene expression assay. Extracted RNA was stored at - 80°C, as well as dilutions of 1 ng/µl nucleic acid for use in multiplex digital polymerase chain reaction (dPCR) analysis.

MDR pump genes *acrA, mdtE,* and *emrA* were probed and assigned a fluorescent channel in the QuantStudio Absolute Q Digital PCR System: FAM (550 nm) - *mdtE*, HEX (555 nm) - *acrA*, and TAMRA (583 nm) - *emrA* (67–70). We used 4X RT Master Mix for RT-PCR reactions, 5X No-RT Master Mix for no-RT reactions (Thermo Fisher Scientific – South San Francisco, CA), and ∼0.9 ng of RNA for each reaction. In the dPCR system, 9 µl of reaction mix was used for the dPCR assay, and the volume was partitioned into >20,000 nano wells. In each nano well, the distribution of templates of each assayed mRNA was assumed to follow the binomial Poisson distribution (71). The optimized dPCR protocol included the following cycling routine: 50°C/10’; 95°C/5’; (95°C/5”; 55°C/30”) x 45 cycles. For each extracted RNA sample, a no-reverse-transcription (no-RT) PCR reaction was performed parallel to the reverse-transcription (RT) reaction in dPCR to check for DNA contamination.

### Statistical Analysis

Mann-Whitney U tests were performed to compare flow cytometry assay selection rates to their respective pH 5.5 control (*p<0.05, **p<0.01, ***p<0.001). Welch’s t-tests were performed to compare cytoplasmic pH assay results to the respective pH 5.5 control (*p<0.05, **p<0.01, ***p<0.001). Mosaic R package was used for Mann–Whitney U tests, Welch’s two-sample t-tests, summary statistics, and Pearson r^2^-values. Mann-Whitney U tests were used for competition assay data because they are not normally distributed. For all figures, statistical test results are presented in **Table S1**.

dPCR analysis was performed in R Studio. The template concentrations (copies/µl) of the three drug pump genes (*acrA*, *mdtE*, *emrA*) were normalized to total RNA of sample. The template concentrations were calculated by the QuantStudio data analysis program based on a Poisson distribution (71). Two-way ANOVA and post-hoc least squares difference with the Benjamini-Hochberg correction were used to compare expression levels of individual efflux pump genes across all tested conditions. Mosaic and Agricolae R packages were used for ANOVA and multiple comparisons.

## AUTHOR CONTRIBUTIONS

YL and AMVH led the team on competition assays and drafted the full manuscript. MTNP devised the dPCR assay and led the expression analysis team, and contributed to the manuscript. BNND, RC, SDRR, KT, and AS performed competition assays. AP and AMVH performed dPCR assays. CCM performed competition assays and trained students. ZCS mentored students and performed pHluorin and competition assays. JLS conceived the study, mentored and supported students, and directed writing of the manuscript.

## ACKNOWLEDGEMENTS

This work was supported by the National Science Foundation awards MCB 1923077 and MRI 1725426. We thank Luke Smallwood for scripting the pHluorin analysis.

