## Supplemental Tables 1-3 for "Fitness Tradeoffs of Multidrug Efflux Pumps in *Escherichia coli* K-12 in Acid or Base, and with Aromatic Phytochemicals"

#### Table S1. Statistical tests of significance for Figures 2 - 7

##### Figure 2A. Mann-Whitney U Test for Difference Between Treatments and pH 5.5 Control

| Condition | Mean | Median | sd | IQR | n | W | p-value | Annotation (*, p < 0.05; **, p < 0.01; ***, p < 0.001) |
|---|---|---|---|---|---|---|---|---|
| *ΔacrA* pH 5.5 Control | 0.28 | 0.402 | 0.639 | 1.09 | 12 | N/A | N/A | N/A |
| *ΔacrA* pH 8.0 Control | -1.34 | -1.27 | 0.413 | 0.337 | 12 | 144 | 7.40E-07 | *** |
| *ΔacrA* pH 5.5 10 µM CCCP | -0.959 | -0.942 | 0.585 | 0.642 | 12 | 133 | 0.000144 | *** |
| *ΔmdtE* pH 5.5 Control | -0.557 | -0.623 | 0.396 | 0.511 | 12 | N/A | N/A | N/A |
| *ΔmdtE* pH 8.0 Control | -1.7 | -1.61 | 0.448 | 0.635 | 12 | 143 | 1.48E-06 | *** |
| *ΔmdtE* pH 5.5 10 µM CCCP | -1.62 | -1.73 | 0.745 | 0.754 | 12 | 131 | 0.000274 | *** |
| *ΔemrA* pH 5.5 Control | -0.327 | -0.374 | 0.208 | 0.274 | 12 | N/A | N/A | N/A |
| *ΔemrA* pH 8.0 Control | -1.18 | -1.18 | 0.126 | 0.204 | 12 | 144 | 7.40E-07 | *** |
| *ΔemrA* pH 5.5 10 µM CCCP | 0.43 | 0.085 | 1.27 | 1.5 | 20 | 61 | 2.11E-02 | * |

##### Figure 2B. Welch's Two Sample t-test for Difference Between Treatments and pH 5.5 Control

| Condition | Mean | Median | sd | IQR | n | t | p-value | Annotation (*, p < 0.05; **, p < 0.01; ***, p < 0.001) |
|---|---|---|---|---|---|---|---|---|
| pH 5.5 Control | 7.21 | 7.18 | 0.203 | 0.317 | 79 | N/A | N/A | N/A |
| 10 µM CCCP | 7.04 | 7.01 | 0.0725 | 0.0799 | 7 | -4.75 | 1.98E-04 | *** |
| 100 µM CCCP | 5 | 5.02 | 0.087 | 0.0401 | 6 | -52.3 | 2.01E-13 | *** |

##### Figure 3. Welch's Two Sample t-test for Difference Between Treatments and pH 5.5 Control

| Condition | Mean | Median | sd | IQR | n | t | p-value | Annotation (*, p < 0.05; **, p < 0.01; ***, p < 0.001) |
|---|---|---|---|---|---|---|---|---|
| pH 5.5 Control | 7.21 | 7.18 | 0.203 | 0.317 | 79 | N/A | N/A | N/A |
| Salicyl-OH | 7.49 | 7.49 | 0.0365 | 0.0564 | 6 | -10.2 | 6.60E-13 | *** |
| Butyrate | 7.04 | 6.97 | 0.151 | 0.234 | 6 | -2.7 | 0.0328 | * |
| Benzyl-OH | 7.42 | 7.56 | 0.275 | 0.388 | 6 | 1.76 | 0.133 | |
| Vanillin | 7.4 | 7.42 | 0.149 | 0.222 | 7 | -3.07 | 0.015 | * |
| Salicylamide | 7.44 | 7.48 | 0.159 | 0.234 | 11 | -4.16 | 0.000839 | *** |
| Sorbate | 6.8 | 6.8 | 0.136 | 0.192 | 11 | 8.78 | 1.02E-07 | *** |
| Ferulate | 7.4 | 7.42 | 0.145 | 0.177 | 7 | 3.06 | 0.015 | * |
| Benzoate | 6.88 | 6.92 | 0.12 | 0.147 | 8 | -6.93 | 1.94E-05 | *** |
| Salicylate | 6.38 | 6.35 | 0.0653 | 0.0561 | 18 | 30.4 | 2.20E-16 | *** |
| Me-Salicylate | 7.3 | 7.01 | 0.175 | 0.124 | 9 | 1.36 | 0.201 | |

**Figure 4. Mann-Whitney U Test for Difference Between Treatments and pH 5.5 Control**

| Condition | Mean | Median | sd | IQR | n | W | p-value | Annotation (*, p < 0.05; **, p < 0.01; ***, p < 0.001) |
|---|---|---|---|---|---|---|---|---|
| pH 5.5 Control | 0.28 | 0.402 | 0.639 | 1.09 | 12 | N/A | N/A | N/A |
| Salicyl-OH | -0.0861 | -0.212 | 0.548 | 0.577 | 12 | 98 | 1.43E-01 | *** |
| Butyrate | -0.15 | -0.0661 | 0.294 | 0.476 | 12 | 105 | 0.0597 | |
| Benzyl-OH | -0.557 | -0.623 | 0.396 | 0.511 | 12 | 123 | 0.00232 | ** |
| Vanillin | 0.193 | 0.0975 | 0.53 | 0.658 | 20 | 138 | 0.502 | |
| Salicylamide | -0.441 | -0.325 | 0.588 | 0.926 | 12 | 116 | 0.01 | * |
| Sorbate | -0.327 | -0.374 | 0.208 | 0.274 | 12 | 113 | 1.73E-02 | *** |
| Ferulate | -0.733 | -0.711 | 0.355 | 0.56 | 12 | 130 | 0.000371 | *** |
| Benzoate | -0.413 | -0.372 | 0.3 | 0.428 | 12 | 115 | 1.21E-02 | *** |
| Salicylate | -1.27 | -1.32 | 0.726 | 0.57 | 12 | 134 | 1.03E-04 | *** |
| Me-Salicylate | -1.54 | -1.61 | 0.481 | 0.791 | 12 | 143 | 1.48E-06 | *** |

**Figure 5. Mann-Whitney U Test for Difference Between Treatments and pH 5.5 Control**

| Condition | Mean | Median | sd | IQR | n | W | p-value | Annotation (*, p < 0.05; **, p < 0.01; ***, p < 0.001) |
|---|---|---|---|---|---|---|---|---|
| pH 5.5 Control | -0.712 | -0.655 | 0.295 | 0.455 | 12 | N/A | N/A | N/A |
| Salicyl-OH | -0.689 | -0.761 | 0.459 | 0.612 | 12 | 76 | 8.43E-01 | |
| Butyrate | -0.638 | -0.536 | 0.374 | 0.597 | 12 | 59 | 0.478 | |
| Benzyl-OH | -0.811 | -0.777 | 0.411 | 0.707 | 12 | 86 | 0.443 | |
| Vanillin | -0.545 | -0.506 | 0.6 | 1.18 | 12 | 64 | 0.671 | |
| Salicylamide | -0.841 | -0.865 | 0.526 | 0.794 | 12 | 88 | 0.378 | |
| Sorbate | -0.567 | -0.532 | 0.368 | 0.666 | 12 | 52 | 2.66E-01 | |
| Ferulate | -0.789 | -0.908 | 0.631 | 0.583 | 12 | 97 | 0.16 | |
| Benzoate | -0.341 | -0.26 | 0.239 | 0.361 | 12 | 27 | 8.29E-03 | * |
| Salicylate | -0.156 | -0.0635 | 0.343 | 0.297 | 12 | 14 | 3.71E-04 | ** |
| Me-Salicylate | -0.895 | -1.02 | 0.497 | 0.826 | 12 | 90 | 3.19E-01 | |

**Figure 6. Mann-Whitney U Test for Difference Between Treatments and pH 5.5 Control**

| Condition | Mean | Median | sd | IQR | n | W | p-value | Annotation (*, p < 0.05; **, p < 0.01; ***, p < 0.001) |
|---|---|---|---|---|---|---|---|---|
| pH 5.5 Control | -0.381 | -0.437 | 0.241 | 0.428 | 12 | N/A | N/A | N/A |
| Salicyl-OH | -0.572 | -0.726 | 0.385 | 0.237 | 12 | 107 | 4.49E-02 | * |
| Butyrate | -0.206 | -0.261 | 0.372 | 0.557 | 12 | 54 | 0.319 | |
| Benzyl-OH | -0.364 | -0.452 | 0.246 | 0.436 | 12 | 72 | 1 | |
| Vanillin | -0.474 | -0.425 | 0.372 | 0.543 | 12 | 86 | 0.443 | |
| Salicylamide | -0.713 | -0.716 | 0.331 | 0.56 | 12 | 110 | 0.0284 | * |
| Sorbate | -0.551 | -0.509 | 0.219 | 0.123 | 12 | 96 | 1.78E-01 | |
| Ferulate | -0.693 | -0.658 | 0.336 | 0.601 | 12 | 106 | 0.0519 | |
| Benzoate | -0.179 | -0.188 | 0.194 | 0.304 | 16 | 46 | 1.98E-02 | * |
| Salicylate | 0.276 | 0.267 | 0.26 | 0.298 | 16 | 4 | 7.89E-07 | *** |
| Me-Salicylate | -0.642 | -0.632 | 0.452 | 0.592 | 12 | 102 | 8.87E-02 | |

**Figure 7. Two-Way ANOVA with Post-Hoc Least Squares Difference Using Benjamini-Hochberg Correction for Gene Expression Differences Between Conditions**

| ***acrA*** | **df** | **Sum of Sqaures** | **Mean of Squares** | **F** | **p-value** |
|---|---|---|---|---|---|
| **Condition** | 6 | 228000 | 38000 | 6.29 | 0.00028 |
| **Growth Phase** | 1 | 336000 | 336000 | 55.6 | 4E-08 |
| **Interaction (Condition:Growth Phase)** | 6 | 545000 | 90800 | 15.0 | 1.28E-07 |
| **Residuals** | 28 | 169000 | 6050 | | |

| ***acrA*** | **Mean** | **Median** | **sd** | **IQR** | **n** | **Multiple Comparisons Results #** |
|---|---|---|---|---|---|---|
| pH 5.5--Log | 372 | 384 | 113 | 112 | 3 | b |
| pH 5.5--Transition | 124 | 130 | 22.9 | 22.2 | 3 | de |
| pH 6.8--Log | 373 | 336 | 66.2 | 57.9 | 3 | b |
| pH 6.8--Transition | 116 | 105 | 26.3 | 24.4 | 3 | de |
| pH 8.0--Log | 305 | 313 | 46.7 | 46.1 | 3 | bc |
| pH 8.0--Transition | 103 | 108 | 10.1 | 9.2 | 3 | de |
| Me-Salicylate--Log | 688 | 718 | 60.6 | 54.7 | 3 | a |
| Me-Salicylate--Transition | 53.4 | 51 | 7.33 | 7.04 | 3 | e |
| Sorbate--Log | 143 | 149 | 16 | 15.2 | 3 | de |
| Sorbate--Transition | 76.4 | 81.7 | 12.3 | 11.4 | 3 | de |
| Benzoate--Log | 372 | 384 | 113 | 30.7 | 3 | cde |
| Benzoate--Transition | 227 | 178 | 119 | 111 | 3 | bcd |
| Salicylate--Log | 314 | 321 | 125 | 147 | 3 | bcd |
| Salicylate--Transition | 224 | 125 | 170 | 125 | 3 | bc |

### Results of post-hoc least squares difference using Benjamini-Hochberg Correction.
Letters indicate groups that are significantly different ($p<0.05$).
Groups that are significantly different do not share a letter and are marked with * in Fig. 7.

| ***mdtE*** | **df** | **Sum of Sqaures** | **Mean of Squares** | **F** | **p-value** |
|---|---|---|---|---|---|
| **Condition** | 6 | 2600000 | 434000 | 116 | 2.20E-16 |
| **Growth Phase** | 1 | 285000 | 285000 | 75.9 | 1.85E-09 |
| **Interaction (Condition:Growth Phase)** | 6 | 229000 | 38000 | 10.2 | 4.46E-06 |
| **Residuals** | 28 | 105000 | 3750 | | |

| ***mdtE*** | **Mean** | **Median** | **sd** | **IQR** | **n** | **Multiple Comparisons Results #** |
|---|---|---|---|---|---|---|
| pH 5.5--Log | 149 | 147 | 41.7 | 41.7 | 3 | c |
| pH 5.5--Transition | 296 | 308 | 57.6 | 56.7 | 3 | b |
| pH 6.8--Log | 14.8 | 18.2 | 6.09 | 5.36 | 3 | d |
| pH 6.8--Transition | 299 | 304 | 15.9 | 15 | 3 | b |
| pH 8.0--Log | 34.7 | 34.9 | 7.13 | 7.13 | 3 | d |
| pH 8.0--Transition | 362 | 357 | 27.3 | 27.1 | 3 | b |
| Me-Salicylate--Log | 161 | 170 | 33.5 | 32.6 | 3 | c |
| Me-Salicylate--Transition | 342 | 339 | 6.24 | 5.76 | 3 | b |
| Sorbate--Log | 916 | 904 | 24.2 | 21.7 | 3 | a |
| Sorbate--Transition | 846 | 777 | 181 | 171 | 3 | a |
| Benzoate--Log | 28.6 | 26.1 | 4.94 | 4.44 | 3 | d |
| Benzoate--Transition | 338 | 317 | 97.1 | 95.4 | 3 | b |
| Salicylate--Log | 99.1 | 75.3 | 43.5 | 38.3 | 3 | cd |
| Salicylate--Transition | 74.5 | 80.3 | 22.8 | 22.2 | 3 | cd |

### Results of post-hoc least squares difference using Benjamini-Hochberg Correction.
Letters indicate groups that are significantly different (p<0.05).
Groups that are significantly different do not share a letter and are marked with * in Fig. 7.

| ***emrA*** | **df** | **Sum of Sqaures** | **Mean of Squares** | **F** | **p-value** |
|---|---|---|---|---|---|
| **Condition** | 6 | 92700 | 15500 | 7.85 | 5.13E-05 |
| **Growth Phase** | 1 | 123000 | 123000 | 62.26 | 1.36E-08 |
| **Interaction (Condition:Growth Phase)** | 6 | 89300 | 14900 | 7.56 | 6.94E-05 |
| **Residuals** | 28 | 55100 | 1970 | | |

| ***emrA*** | **Mean** | **Median** | **sd** | **IQR** | **n** | **Multiple Comparisons Results #** |
|---|---|---|---|---|---|---|
| pH 5.5--Log | 176 | 204 | 55.3 | 49.6 | 3 | bc |
| pH 5.5--Transition | 65.4 | 66.7 | 13.5 | 13.5 | 3 | def |
| pH 6.8--Log | 147 | 140 | 29.7 | 29.1 | 3 | bcd |
| pH 6.8--Transition | 22.6 | 20.7 | 5.99 | 5.75 | 3 | f |
| pH 8.0--Log | 74.1 | 81.5 | 13.2 | 11.6 | 3 | def |
| pH 8.0--Transition | 9.43 | 8.63 | 1.79 | 1.65 | 3 | f |
| Me-Salicylate--Log | 295 | 310 | 53.5 | 51.9 | 3 | a |
| Me-Salicylate--Transition | 11.9 | 11.6 | 0.582 | 0.536 | 3 | f |
| Sorbate--Log | 203 | 195 | 16.8 | 15.5 | 3 | b |
| Sorbate--Transition | 50.8 | 47.1 | 9.09 | 8.52 | 3 | ef |
| Benzoate--Log | 173 | 194 | 39.3 | 34.8 | 3 | bc |
| Benzoate--Transition | 107 | 53.1 | 99.6 | 87.9 | 3 | cde |
| Salicylate--Log | 180 | 145 | 68.4 | 61.5 | 3 | bc |
| Salicylate--Transition | 224 | 205 | 62.1 | 60.1 | 3 | ab |

### Results of post-hoc least squares difference using Benjamini-Hochberg Correction.
Letters indicate groups that are significantly different (p<0.05).
Groups that are significantly different do not share a letter and are marked with * in Fig. 7.

**Table S2. Culture times and $OD_{600}$ values at harvest for RNA preparation**

| Media | Log-phase $OD_{600}$ | Log-phase growth time | Transition-phase $OD_{600}$ | Transition-phase growth time |
|---|---|---|---|---|
| **pH 5.5 Control** | 0.2, 0.15, 0.14 | 3 h | 0.44, 045, 0.43 | 7 h |
| **pH 6.8 Control** | 0.12, 0.12, 0.11 | 3 h | 0.73, 0.66, 0.69 | 7 h 15 min |
| **pH 8.0 Control** | 0.18, 0.16, 0.15 | 3 h | 0.56, 0.57, 0.56 | 7 h 15 min |
| **pH 5.5 1 mM Methyl salicylate** | 0.11, 0.11, 0.12 | 3 h 40 min | 0.51, 0.50, 0.50 | 11 h 20 min |
| **pH 5.5 1 mM Sorbate** | 0.17, 0.16, 0.17 | 5 h | 0.64, 0.53, 0.58 | 7 h 30 min |
| **pH 5.5 1 mM Benzoate** | 0.13, 0.14, 0.13 | 5 h 20 min | 0.42, 0.38, 0.44 | 7 h 50 min |
| **pH 5.5 1 mM Salicylate** | 0.15, 0.16, 0.17 | 6 h 30 min | 0.39, 0.44, 0.52 | 11 h |

**Table S3: Taqman probes and primers used to specifically amplify *mdtE, acrA, emrA,* and *rpoB* cDNA in 4-plex dPCR gene expression assay.** Probes and primers were designed by IDT (Coralville, IA).

| Gene | Forward Primer | Reverse Primer | Taqman Probe* | Amplicon Length |
|---|---|---|---|---|
| ***acrA*** | 5'-CAGTAAGCAAGAGTACGATCAGG | 5'-GGAGAGGTGACTTTGGTGTAAG | 5'- HEX-CAACAGGCGAATGCTGCGGTAAC | 123bp |
| ***mdtE*** | 5'-CATGTACGTCACGGCATTAGT | 5'-TGCACGACATCGTCTTTATCC | 5'- 6-FAM-ACATTCTGGCGGCTACCTTCATCC | 129bp |
| ***emrA*** | 5'-GCAAATGCGGAGACTCAAAC | 5'-CCCTATCGCTACGGCAATAAT | 5'- 6-TAMRA-AAAGAGCAAGGTGAGAAGGAGGAGC | 111bp |
| ***rpoB*** | 5'-TTCTTCTCCGAAGACCGTTATG | 5'-GATACCGGAACCTTCGATTTCT | 5'- TYE665-TGTCTGCGGTTGGTCGTATGAAGT | 93bp |

*****Fluorophores: HEX, hexachlorofluorescein; 6-FAM, fluorescein; 6-TAMRA, 6-Carboxytetramethylrhodamine; TYE665, proprietary IDT
